# Protein Design by Directed Evolution Guided by Large Language Models

**DOI:** 10.1101/2023.11.28.568945

**Authors:** Thanh V. T. Tran, Truong Son Hy

## Abstract

Directed evolution, a strategy for protein engineering, optimizes protein properties (i.e., fitness) by a rigorous and resource-intensive process of screening or selecting among a vast range of mutations. By conducting an *in silico* screening of sequence properties, machine learning-guided directed evolution (MLDE) can expedite the optimization process and alleviate the experimental workload. In this work, we propose a general MLDE framework in which we apply recent advancements of Deep Learning in protein representation learning and protein property prediction to accelerate the searching and optimization processes. In particular, we introduce an optimization pipeline that utilizes Large Language Models (LLMs) to pinpoint the mutation hotspots in the sequence and then suggest replacements to improve the overall fitness. Our experiments have shown the superior efficiency and efficacy of our proposed framework in the conditional protein generation, in comparision with other state-of-the-art baseline algorithms. We expect this work will shed a new light on not only protein engineering but also on solving combinatorial problems using data-driven methods. Our implementation is publicly available at https://github.com/HySonLab/Directed_Evolution.

## I. Introduction

**P**ROTEIN are essential biomolecules that play a diverse range of critical roles in every living organism. They are involved in virtually every important activity that happens inside living things: digesting food, contracting muscles, moving oxygen throughout the body, attacking foreign viruses, etc. The effective resolutions to biological challenges can also serve as effective resolutions to human challenges, given the extensive utilization of proteins in various domains such as food, chemicals, consumer products, and medicinal applications. However, comprehending the complex structure and function of proteins remains an immensely challenging task, owing to the diversity of protein sequences and the three-dimensional structure they have. Therefore, computational biology and machine learning (ML) methods have emerged as indispensable tools among researchers and engineers. These methods offer the capability to process vast amounts of biological data, extract meaningful patterns, and make predictions that would be impractical through traditional experimental approaches alone [1]. Moreover, large language models have been pretrained on extensive protein sequence databases [2], [3], enabling them to capture intricate sequence-structure-function relationships. These models empower researchers to extract valuable insights from protein sequences, understand their evolution, and predict their behavior in various biological contexts.

In the field of protein engineering, fitness can be defined as performance on a desired property or function. Examples of fitness include catalytic activity for enzymes [4] and fluorescence for biomarkers [5]. Protein optimization seeks to improve protein fitness by altering the underlying sequences of amino acids. Nevertheless, the space of possible proteins is too large to search exhaustively naturally, in the laboratory, or computationally. The problem of the identification of ideal sequences is classified as NP-hard, as there currently exists no polynomial-time algorithm for efficiently exploring this particular domain [6]. Functional proteins are scarce in this vast space of sequences, and as the desired level of function increases, the number of sequences having that function decreases exponentially [7]. Consequently, the occurrence of functional sequences is rare and overshadowed by nonfunctional and mediocre sequences. In recent years, generative models have emerged as powerful tools for generating creative and diverse content, ranging from text [8], [9], images [10], [11] to speech [12] and more. Leveraging the success of generative models in text and image generation, numerous studies have been made to harness these techniques to design novel proteins with tailored properties and functions [13]– [15]. This developing paradigm promises to accelerate wet-lab experiments with data-driven models to find desirable proteins more efficiently.

Directed evolution, a laboratory methodology wherein biological entities possessing desired characteristics are generated through iterative cycles of genetic diversification and library screening or selection, has emerged as a highly valuable and extensively utilized instrument in both fundamental and practical realms of biological research [16]–[19]. This method was conceptualized using the natural process of evolution and the idea of natural selection as its primary sources of inspiration. In nature, organisms with advantageous traits are more likely to survive and reproduce, passing on their beneficial characteristics to the next generation. Similarly, directed evolution starts with a protein having some level of the desired function. Subsequently, a series of mutation and screening rounds are undertaken, wherein mutations are introduced (e.g., through site-saturation mutagenesis) to generate a collection of variant proteins. Through this iterative process, the most optimal variant is identified, and the cycle continues until a satisfactory level of improvement is achieved. Fig. 1A demonstrates the workflow of this algorithm.

**Fig. 1.**
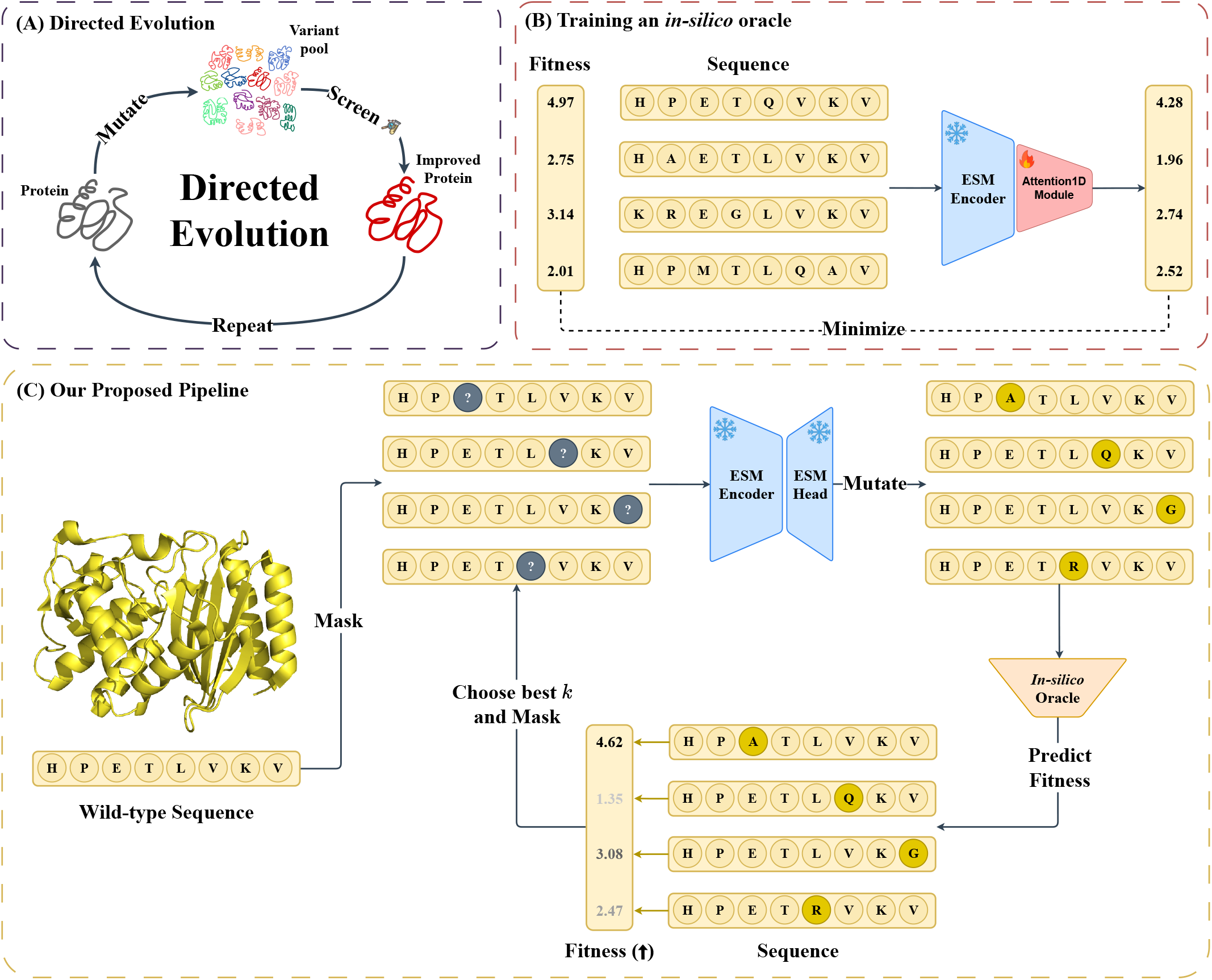
**(A)** Workflow of traditional directed evolution: many variants are screened, and the best variant is selected as the parent for the next round of mutagenesis and screening. **(B)** We train the *in-silico* oracle as the “ground-truth” evaluator to predict the fitness score of each generated sequence. **(C)** Our proposed MLDE framework. The workflow begins by identifying a protein with activity for a target function. Once the starting point is identified, diversity is introduced through mutagenesis, and the fitness scores of the resulting variants are predicted by a trained *in-silico* oracle. Based on these predictions, we can select multiple variants as potential parent candidates for the subsequent iteration, balancing exploitation and exploration.

Machine learning-guided directed evolution (MLDE) is a strategy for protein engineering that can be applied to a range of biological systems, reducing the burden of laboratory experiments by performing *in-silico* population selection through model-based fitness prediction. These techniques can leverage data from both improved and unimproved sequences in order to accelerate the process of evolution and broaden the range of properties that can be optimized. This is achieved by intelligently selecting new variants for screening, which enables the achievement of higher levels of fitness than what can be accomplished solely through directed evolution [20]– [22]. Employing this novel biology algorithm, we propose a deep learning-based framework for MLDE that optimizes directly on the discrete space of all possible sequences. Our pipeline can be described by Fig. 1C.

Our contributions can be summarized as follows:

- We propose a novel LLM-guided mutation prediction paradigm framework for *in-silico* directed evolution,
- We introduce a masking strategy that aids the identification of promising mutations,

## II. Related work

### A. Directed Evolution

Directed Evolution is a classical paradigm for protein sequence design that has achieved several successes. Under this framework, ML algorithms play a crucial role in improving the sample efficiency of evolutionary search. [23] utilizes a fast Markov chain Monte Carlo sampler that uses gradients to propose promising mutations. [24] formulates the local search as a proximal optimization problem to derive an objective for searching the steepest improvement. [25] combines hierarchical unsupervised clustering sampling and supervised learning to guide protein engineering. This work is later improved by [26]. [27] provides an approach to target informative sequences without the initial global search. Nevertheless, these methods have not yet incorporated large language models, which have proven efficient in multiple related tasks. Our proposal describes the combination of a protein language model yielding a highly favorable outcome.

### B. Protein Property Prediction

One of the primary objectives of bioinformatics is the prediction of protein function, which can be applied to a vast array of biological issues, including the identification of drug targets and the comprehension of disease mechanisms. Numerous computational approaches have been developed to autonomously forecast protein function, with specific investigations concentrating on site- or domain-specific predictions [28], [29]. Traditional machine learning classifiers, such as support vector machines, random forests, and logistic regression, have been extensively used for protein function prediction [30], [31]. In recent years, deep learning has led to unprecedented improvements in the performance of methods tackling a broad spectrum of problems, including predicting protein function [32]–[37]. These developments facilitate our work as we employ attention, a well-known mechanism in deep learning, to acquire a comprehensive understanding of the protein sequence and anticipate its properties.

### C. Protein Generation

Significant advancements have been made in the development of methodologies for the generation of functional protein sequences with desired features in recent times. Conventional approaches that utilize multiple sequence alignments of homologous proteins, such as ancestral sequence reconstruction [38], have exhibited efficacy in producing functional proteins, but with certain limitations in their applicability. To access a broader sequence space, different ML techniques have been employed. Some techniques, including reinforcement learning [39], [40], Bayesian optimization [41]–[43], importance sampling [44], and adaptive evolution method [45] have been used and achieved great results in their defined tasks. Modern algorithms such as language models provide a powerful architecture to learn from large sets of amino acid sequences across families for the purpose of generating diverse, realistic proteins [46]–[48]. Other generative techniques like variational auto-encoder [49], [50], generative adversarial networks [14], [51], and diffusion [52] also show promising outcomes in numerous protein engineering tasks. Yet, by applying MLDE to generating sequences, we find that this algorithm demonstrates strong performance, especially in comparison to other state-of-the-art (SOTA) methods.

## III. Background

### A. Notation and Problem Formulation

Through the process of directed evolution, protein fitness is enhanced by emulating natural selection through mutagenesis. From a mathematical standpoint, directed evolution can be conceptualized as an optimization problem for a black-box algorithm, seeking the optimal sequence *x*^⋆^

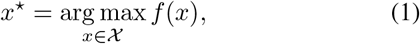

where 𝒳 is the sequence mutation space, and *f* (*x*) is an unknown sequence-to-fitness function for sequence *x* in 𝒳. During the optimization, the algorithm choose the best mutated sequence from an intermediate generated population **x** (the size of the population |**x**| is *n*) so that the returned fitness *y*^⋆^ = *f* (*x*^⋆^) is the best fitness among values **y**.

### B. Transformer

Transformer [53] is a formidable force in the modern deep learning stack. Originally, Transformer follows an encoder-decoder scheme and employs a multi-headed self-attention mechanism. The fundamental concept underlying this mechanism is for each element within the sequence to acquire information from other tokens present in the same sequence. Concretely, given a tensor of sequence features **X** ∈ ℝ^*n×d*^, Transformer computes three matrices including query (**Q**), key (**K**), and value (**V**) via three linear transformations 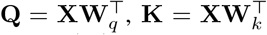, and 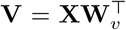. A self-attention tensor (**H**) can be computed as follows:

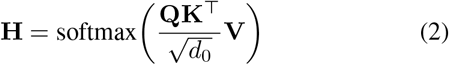

where **W**_*q*_, **W**_*k*_, and **W**_*v*_ are learnable parameters in 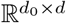, resulting in 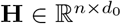. Moreover, each **H** in Eq. 2 denotes an attention head. The output of multiple heads (**H**_1_, … , **H**_*h*_) are then concatenated together to form a final tensor **H** = Concat(**H**_1_, … , **H**_*h*_), where *h* is the number of attention heads. Finally, the new representation **X**^*′*^ can be computed by feeding **H** into a feed-forward neural network (FFN).

### C. ESM-2: Protein Language Model

Inspired by [54], ESM-2 [55] adopts the encoder-only Transformer architecture style with small modifications. The original Transformer uses absolute sinusoidal positional encoding to inform the model about token positions. These positional encodings are added to the input embeddings at the bottom of the encoder stack. Nevertheless, this approach demonstrates limited generalizability beyond the specific context window on which it is trained. The implementation of Rotary Positional Embedding (RoPE) [56] in ESM-2 enables the model to extrapolate beyond its trained context window with no additional computational effort. The ESM-2 model is capable of generating latent representations for individual amino acids inside a protein sequence. This is achieved through pre-training on a vast dataset consisting of millions of protein sequences including billions of amino acids. Specifically, given an input protein, 15% of amino acids are masked and ESM-2 is tasked with predicting these missing positions. Although this training objective only directly involves predicting missing amino acids, achieving a high degree of success requires the model to learn complex internal representations of its input. Consequently, the model is able to produce a concise representation of the full protein sequence, without relying on three-dimensional information. This rich and meaningful representations of ESM-2 has aided numerous studies, including protein functional prediction [57], [58], protein structure prediction [55], protein-protein interaction prediction [59], protein multimodal representation [60], and protein design [61], [62].

## IV. Method

This section describes the overall pipeline of our framework. Initially, after dividing the sequence population into multiple parts, we apply different masking strategies for each portion to retrieve a partially-masked population. These masked variants are then fed into ESM-2 to predict the tokens being masked. After that, the newly generated population is measured for fitness by a fine-tuned Attention1D. These steps are repeatedly iterated until they reach a pre-defined iteration number. The full workflow of our method is illustrated in Fig. 1C.

### A. Selecting Mutation Positions

We follow the scheme of the text-based masked language model [54], which masks some of the tokens from the input and predicts the original vocabulary id of the masked. In the context of protein generation, we assume that the new prediction tokens can boost the fitness values of the protein sequence. In this study, we conducted two different masking strategies: **(1)** random masking and **(2)** importance masking. In the first scheme, we randomly replace token(s) in the sequence by the ⟨`MASK`⟩ tokens. For the latter, we follow the softmasking strategy described in [63]. Concretely, when we split a protein sequence into multiple substrings of length *k* called *k-mers*, it creates a distribution based on the frequency of each k-mer. Therefore, when masking and recovering the sequences, k-mer with various importance should be treated differently. Following [63], we consider the following aspects to measure and define the importance of k-mers in one sequence:

#### 1) Relevancy

We adopt the TF-IDF weights [64], *w*_TF-IDF_, of the k-mer as one way to measure the relevance of k-mer *w* in one sequence *x*:

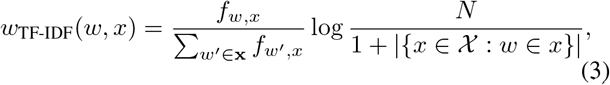

where *f*_*w*,*x*_ is the number of times that k-mer *w* occurs in sequence **x**, *N* is the number of sequences in the corpus, and 𝒳 is the set of sentences, and |{*x 𝒳*∈ : *w* ∈ *x*}| is the number of sequences where the k-mer *w* appears. A higher weight for k-mer *w* in sequence *x* in TF-IDF means that the k-mer might be more important in the sequence.

#### 2) Entropy

We also consider measuring the amount of information with entropy *H* [65] in the k-mer *w* to reflect the importance of that k-mer:

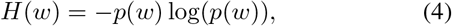

where 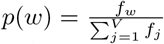 represents the probability of k-mer *w* and *f* is the number of appearances of k-mer in the corpus. A k-mer with lower entropy indicates that it might contain less information and thus be less important compared to the one with higher entropy.

In practice, we combine these two measures (with normalization) to decide the importance *I* of the k-mer *w* in one sequence *x* by:

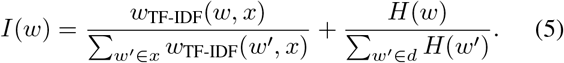

Based on the introduced importance *I* of the k-mers in a sequence, we decide to replace k-mer(s) with the lowest importance value(s) by ⟨`MASK`⟩ token(s). By doing this, models could replace the least important k-mers with better ones for better generation quality.

### B. Machine Learning-guided Improvement

#### Algorithm 1

Machine Learning-guided Directed Evolution

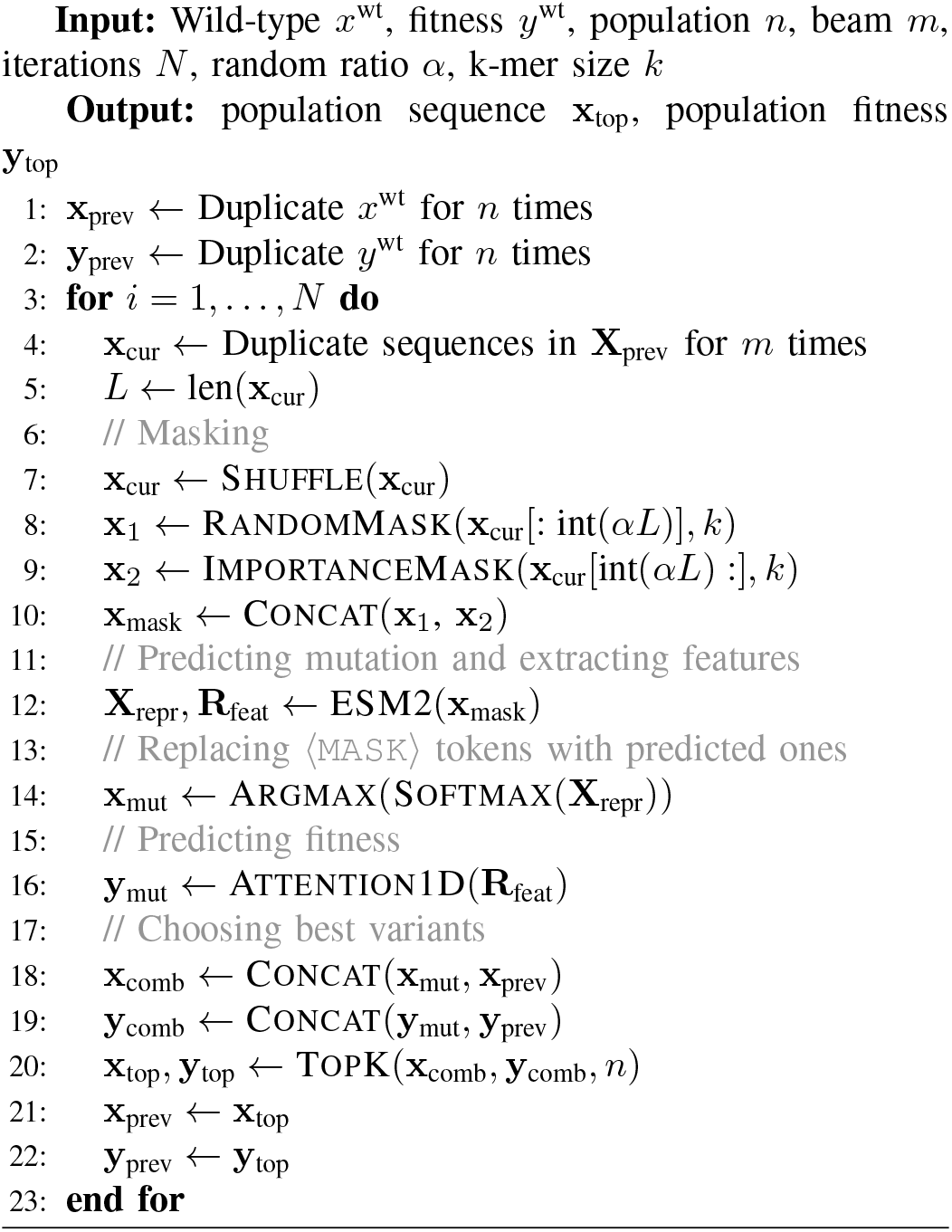

After the generation of a population of masked sequences, these sequences are subsequently inputted into the ESM-2 model. Following this, the neural network generates predictions for the masked tokens and produces a latent representation of these variations. In the final stage, we evaluate the sequences that have been newly proposed by measuring their fitness values. In practice, validating these generated sequences experimentally is costly and time-intensive. We follow prior works [24], [44] in leveraging a trained evaluator model (i.e., oracle) as a proxy for wet-lab measurements. This oracle operates on the input representation 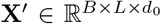 derived from a pre-trained ESM model after processing a protein sequence. The architecture of this module is as follows:

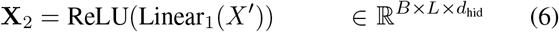

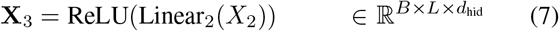

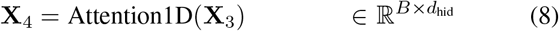

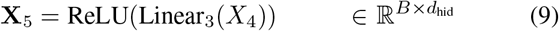

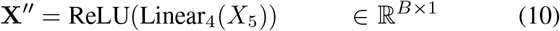

where *B* is the number of sequences, *L* is the input length, and *d*_hid_ is the hidden dimension of the module. As illustrated in the Figure 1B, this model is subsequently trained to comprehend the fitness landscape of the specified protein sequence. The values are thereafter arranged in descending order to generate the most optimal sequences that will proceed to the subsequent iteration of the pipeline.

In summary, our pipeline can be described by Algorithm 1. It is important to note that while selecting the best variants in each iteration, we combine the present “effective” population with the population from the preceding iteration. This procedure guarantees that in the event of mutations resulting in a reduction in fitness values, the pipeline will opt for the variations from the preceding round rather than the inferior ones.

## V. Experiments

In this section, we conduct extensive experiments to validate the effectiveness of our proposed pipeline on protein sequence design task.

### A. Datasets

Following [24] and [44], we evaluate our method on the following eight protein engineering benchmark datasets: (1) Green Fluorescent Protein (**avGFP**), (2) Adeno-Associated Viruses (**AAV**), (3) TEM-1 *β*-Lactamase (**TEM**), (4) Ubiquitination Factor Ube4b (**E4B**), (5) Aliphatic Amide Hydrolase (**AMIE**), (6) Levoglucosan Kinase (**LGK**), (7) Poly(A)-binding Protein (**Pab1**), (8) SUMO E2 Conjugase (**UBE2I**). Table I provides comprehensive details of the eight datasets utilized in this study, including the protein name, its organism, sequence length, dataset size, optimization target, and percentile distribution. Each dataset corresponds to different optimization targets, which we simplify as a singular term *“fitness”* when reporting the benchmark results in this paper. The detailed data descriptions are provided in the Supplementary Material.

**Table I.** Detailed information and statistics of the eight protein datasets.

| Dataset | Organism | Protein | Optimization Target | Length | Size | 0.25 | Percentiles<br>0.50 | 0.75 |
| --- | --- | --- | --- | --- | --- | --- | --- | --- |
| avGFP [66] | Aequorea victoria | GFP | Brightness | 237 | 51,715 | 1.428 | 3.287 | 3.161 |
| AAV [67] | Homo sapiens | VP1 | AAV viabilities | 28 | 42,330 | -3.964 | -0.840 | 1.321 |
| TEM [68] | Escherichia coli | TEM-1 β-Lactamase | Thermodynamic stability | 286 | 5,199 | 0.049 | 0.444 | 0.934 |
| E4B [69] | Mus musculus | UBE4B | Ubiquitin ligase activity | 102 | 91,032 | -1.830 | -0.984 | -0.093 |
| AMIE [70] | Escherichia coli | Amidase | Hydrolysis activity | 341 | 6,417 | -1.228 | -0.666 | -0.263 |
| LGK [71] | Lipomyces starkeyi | Levoglucosan kinase | Levoglucosan utilization | 439 | 7,633 | -0.871 | -0.562 | -0.394 |
| Pab1 [72] | Saccharomyces cerevisiae | Poly(A)-binding | mRNA binding | 75 | 36,389 | -0.116 | -0.022 | 0.036 |
| UBE2I [73] | Homo sapiens | UBE2I | Growth rescue rate | 159 | 3,022 | 0.068 | 0.492 | 0.766 |

### B. Implementation Details

Starting with a population of wild-type protein sequences, we split them into two separate groups. Each group is then subjected to a specific masking strategy, resulting in a population of masked sequences. These sequences are then fed to the pretrained 35-million-parameter version of the ESM-2 model, which generates the representation of the sequences and introduces mutations to sequences by replacing the ⟨`MASK`⟩ with the amino acid that the model confidently predicted. This representation is then fed into the oracle described in Section IV-B to predict the fitness value of sequences. Subsequently, the population is arranged in a descending order to select the optimal *n* variations according to their respective fitness values. These variants are the prospective candidates that will undergo the same pipeline in the subsequent iteration. Following several rounds, we would observe enhanced protein sequences exhibiting improved fitness ratings.

In our settings, we set the population size to 128 following [40]. Inspired by the beam search algorithm, which has proven its efficiency in the field of natural language processing [74], [75], the model proposes 4 separate mutated sequences (i.e., a beam size of 4), instead of 1 mutation in traditional directed evolution, to each variant in the population, making the *effective* population size to 512. This adaptation is illustrated in Fig. 2. We configure the random ratio to 0.6 and the k-mer size to 1 for the remaining hyperparameters. It is important to acknowledge that the outcomes generated from these configurations are not guaranteed to be optimal. The examination of the influence of each hyperparameter is conducted in Section V-F.

**Fig. 2.**
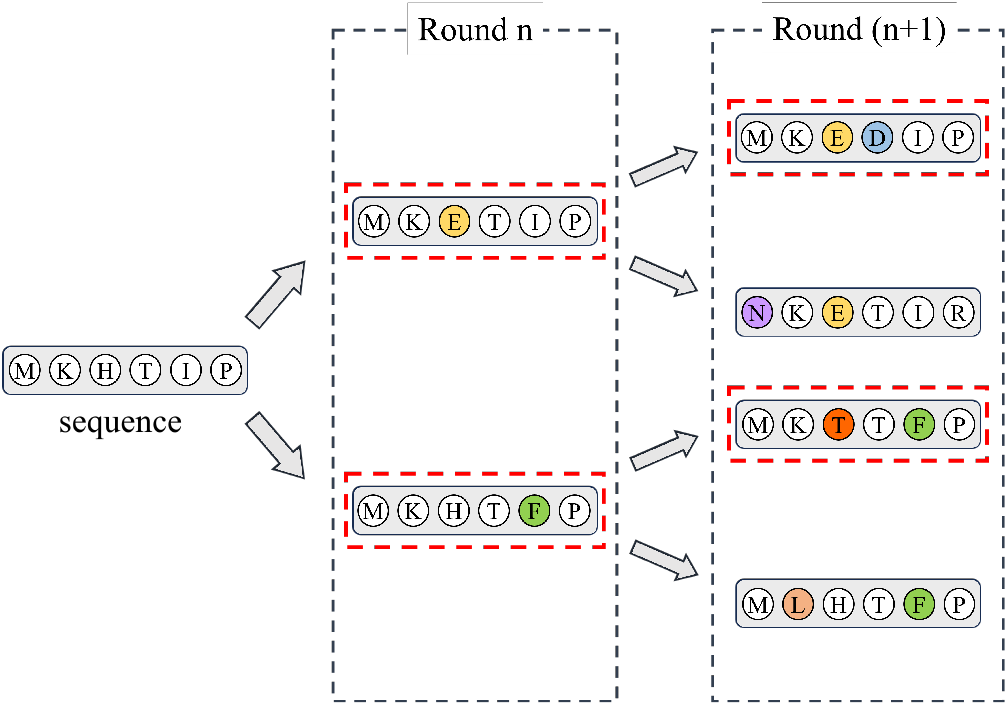
Illustration of beam search adaptation in directed evolution. In this figure, the population number is 2, and the beam size is 2, which makes the *effective* population size 4 (i.e., sequences in the third generation before dropping inferior ones). The sequences covered in red boxes are chosen to move to the next iteration.

To ensure unbiased evaluation and avoid circular use of oracles, which can potentially cause information leakage, we employ two separate oracles for each fitness dataset: (1) the optimization oracle that guides the model optimization and (2) the evaluation oracle that assesses the performance of every methods. Following [24], we construct both oracles by freezing the ESM-based encoders and training an Attention1D module stacked atop to predict fitness scores. We use the pre-trained 33-layer ESM-2 for the former and the trained oracle provided by [24] for the latter. Details on these oracles are provided in the Supplementary Material. For each algorithm, we run all the experiments five times and report the average scores with their standard deviation.

### C. Baseline Algorithms

We assess the performance of our method in comparison to several representative baselines: (1) **AdaLead** [76] is an advanced implementation of model-guided evolution. (2) **DyNA PPO** [39] applies proximal policy optimization to search sequences on a learned landscape model. (3) **CMA-ES** [77] is a famous evolutionary search algorithm. (4) **DbAS** [78] is a probabilistic modeling framework employing an adaptive sampling algorithm. (5) **CbAS** [79] is an improvement over DbAS that conditions on desired properties. (6) **COMS** [80] is conservative objective models for offline MBO. (7) **PEX** [24] is a model-guided sequence design algorithm using proximal exploration. (8) **GFN-AL** [40] applies GFlowNet to design biological sequences. (9) **GGS** [81] is a graph-based smoothing method to optimize protein sequences. To ensure precise evaluation, we re-execute and re-evaluate baseline methods using the same oracle. For implementations from (1) to (5), we utilized the open-source implementation provided by [76]. As for other baseline methods, we adapted and modified the codes released by their respective authors. Details about the hyperparameter selection for these baselines are discussed in the Supplementary Material.

### D. Evaluation Metrics

We employ three metrics from [40] to assess our sequence design pipeline: (1) **MFS**: Maximum fitness score, (2) **Diversity**, and (3) **Novelty**. Additionally, we consider two additional metrics for robustness: (4) **AFS**: Average fitness score, and (5) **dist(WT)**: the distance between the best-designed sequence and wild-type one. Detailed descriptions and mathematical formulations of these metrics can be found in the Supplementary Material. It’s important to note that while greater diversity, novelty, and dist(WT) provide insights into the exploration and exploitation trade-offs, they do not necessarily indicate superior performance.

### E. Experimental Results

#### 1) Comparison with other baselines

As described in Table II, our approach exhibits superior fitness scores in 7 out of 8 protein families when evaluated against methods utilizing the same oracle. Notably, it is observed that AAV and Pab1 contain sequences with limited lengths, up to 28 and 76 amino acids, respectively. However, for datasets featuring longer sequences, leveraging the capabilities of protein language models trained on millions of protein sequences notably enhances MLDE’s optimization capabilities. This is particularly evident on the avGFP and E4B benchmarks, where our method achieves statistically significant outperformance over the baselines. These findings suggest that our approach adeptly harnesses the potential of protein language models to introduce novel mutations into existing proteins.

**Table II.**
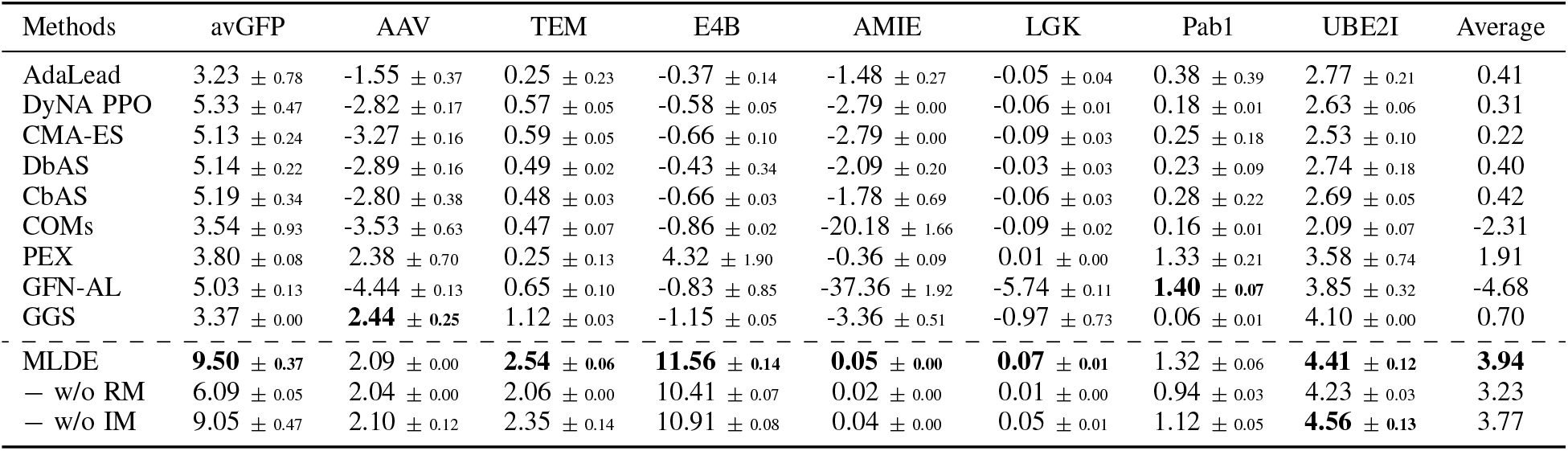
Maximum fitness scores (MFS) of all methods across eight datasets. Higher values indicate better functional properties in the dataset. **B****old results** indicate the best value among all methods assessed using the same oracles.

Additionally, Tables III and IV outline the diversity and novelty metrics for all methods. It is crucial to emphasize that while these results offer insights into the exploration and exploitation trade-off, they may not necessarily reflect the efficacy of an algorithm, as it largely depends on the specific objectives of each method. Some methods aim to optimize the fitness score while maximizing diversity and novelty [44], while others may prioritize minimizing novelty [24], and some may solely focus on optimizing the fitness score alone [81]. Moreover, a random algorithm can achieve maximum diversity and novelty scores. This is evident in the case of GFN-AL and COMs in the AMIE dataset, where it attains high novelty and diversity score, yet achieves the lowest MFS and AFS among all methods.

**Table III.** Diversity of all methods across eight datasets. This table provides insight into the exploration and exploitation trade-off among methods.

| Methods | avGFP | AAV | TEM | E4B | AMIE | LGK | Pab1 | UBE2I | Average |
| --- | --- | --- | --- | --- | --- | --- | --- | --- | --- |
| AdaLead | 9.81 ± 3.23 | 6.90 ± 0.52 | 8.07 ± 4.11 | 6.90 ± 1.56 | 7.43 ± 0.37 | 7.44 ± 3.96 | 7.44 ± 4.13 | 7.42 ± 0.58 | 7.68 |
| DyNA PPO | 204.38 ± 0.14 | 114.23 ± 0.13 | 157.96 ± 0.78 | 140.62 ± 0.15 | 171.12 ± 0.22 | 205.10 ± 1.11 | 185.16 ± 0.09 | 179.21 ± 0.11 | 169.72 |
| CMA-ES | 173.37 ± 4.23 | 20.40 ± 0.58 | 210.14 ± 1.76 | 73.65 ± 0.22 | 248.20 ± 2.97 | 319.89 ± 54.05 | 54.05 ± 1.64 | 116.78 ± 1.34 | 152.06 |
| DbAS | 205.72 ± 0.06 | 25.14 ± 0.03 | 247.72 ± 0.04 | 89.60 ± 0.05 | 294.52 ± 0.06 | 378.22 ± 0.06 | 66.42 ± 0.06 | 138.73 ± 0.07 | 180.69 |
| CbAS | 205.71 ± 0.05 | 25.17 ± 0.02 | 247.63 ± 0.03 | 89.63 ± 0.01 | 294.50 ± 0.08 | 378.31 ± 0.11 | 66.43 ± 0.05 | 138.79 ± 0.08 | 180.77 |
| COMs | 70.32 ± 9.97 | 45.76 ± 0.39 | 73.19 ± 6.49 | 68.40 ± 0.50 | 100.73 ± 1.70 | 126.62 ± 10.61 | 115.24 ± 0.38 | 109.65 ± 2.69 | 88.74 |
| PEX | 7.05 ± 0.97 | 4.78 ± 0.51 | 6.37 ± 0.90 | 6.20 ± 0.94 | 8.00 ± 0.93 | 7.84 ± 1.16 | 7.01 ± 1.95 | 8.27 ± 1.73 | 6.94 |
| GFN-AL | 6.25 ± 0.25 | 0.50 ± 0.02 | 1.23 ± 0.04 | 29.06 ± 0.44 | 46.58 ± 0.22 | 112.75 ± 2.51 | 27.13 ± 0.65 | 2.35 ± 0.07 | 28.23 |
| GGs | 4.63 ± 0.54 | 12.71 ± 0.78 | 7.39 ± 0.67 | 10.58 ± 1.57 | 14.69 ± 0.14 | 16.15 ± 0.25 | 16.73 ± 0.38 | 3.01 ± 0.17 | 10.74 |
| MLDE | 4.92 ± 1.48 | 1.90 ± 0.07 | 2.32 ± 0.85 | 2.32 ± 0.84 | 4.86 ± 1.60 | 6.83 ± 1.20 | 1.69 ± 2.14 | 1.04 ± 0.70 | 3.11 |

**Table IV.** Novelty of all methods across eight datasets. This table provides insight into the exploration and exploitation trade-off among methods.

| Methods | avGFP | AAV | TEM | E4B | AMIE | LGK | Pab1 | UBE2I | Average |
| --- | --- | --- | --- | --- | --- | --- | --- | --- | --- |
| AdaLead | 13.49 ± 5.16 | 17.81 ± 0.48 | 41.41 ± 25.30 | 48.93 ± 2.71 | 41.17 ± 3.94 | 78.87 ± 32.10 | 73.53 ± 20.64 | 78.59 ± 12.63 | 49.23 |
| DyNA PPO | 201.70 ± 0.49 | 111.57 ± 0.10 | 156.78 ± 1.00 | 139.23 ± 0.27 | 170.37 ± 0.32 | 205.82 ± 0.94 | 185.69 ± 0.15 | 179.80 ± 0.27 | 168.87 |
| CMA-ES | 202.11 ± 0.43 | 20.77 ± 0.31 | 247.97 ± 1.08 | 85.57 ± 1.44 | 232.00 ± 9.96 | 375.44 ± 12.31 | 63.95 ± 1.30 | 138.07 ± 0.83 | 170.74 |
| DbAS | 201.80 ± 0.19 | 21.26 ± 0.15 | 247.49 ± 0.37 | 86.29 ± 0.12 | 293.44 ± 0.22 | 382.31 ± 0.37 | 64.93 ± 0.17 | 138.80 ± 0.19 | 179.53 |
| CbAS | 201.79 ± 0.31 | 21.31 ± 0.08 | 247.44 ± 0.35 | 86.13 ± 0.17 | 293.47 ± 0.20 | 382.27 ± 0.07 | 65.06 ± 0.20 | 138.91 ± 0.16 | 179.55 |
| COMs | 184.18 ± 0.35 | 98.83 ± 0.53 | 111.93 ± 1.66 | 101.93 ± 0.32 | 123.59 ± 1.95 | 150.66 ± 1.10 | 134.30 ± 1.97 | 129.80 ± 0.81 | 129.40 |
| PEX | 4.32 ± 0.69 | 1.91 ± 0.23 | 4.58 ± 0.89 | 5.12 ± 0.45 | 4.13 ± 0.66 | 16.35 ± 0.96 | 4.32 ± 1.40 | 5.40 ± 1.61 | 5.77 |
| GFN-AL | 220.63 ± 8.95 | 24.95 ± 1.21 | 255.94 ± 5.06 | 90.63 ± 1.03 | 326.26 ± 4.97 | 413.95 ± 2.08 | 64.05 ± 0.51 | 145.46 ± 1.92 | 192.73 |
| GGs | 3.55 ± 0.27 | 2.06 ± 0.15 | 4.03 ± 0.36 | 8.08 ± 0.33 | 9.99 ± 0.01 | 20.99 ± 0.01 | 8.38 ± 0.10 | 2.86 ± 0.16 | 7.49 |
| MLDE | 103.35 ± 2.11 | 2.58 ± 0.02 | 70.16 ± 2.45 | 28.30 ± 2.63 | 122.83 ± 8.07 | 156.66 ± 5.76 | 23.46 ± 1.26 | 41.02 ± 4.68 | 68.55 |

#### 2) Average performance

Another crucial factor determining the efficacy of an optimization algorithm is its ability to consistently produce high-quality samples within the population. Table V illustrates this aspect through the computation of the AFS, showcasing our method’s prowess as it consistently outperforms all evaluated methods across 8 benchmarks. As a result, our method achieves the top performance, highlighting the highest average score of 3.669, surpassing the runner-up, PEX, by a significant margin of 5.6-fold.

**Table V.** Average fitness scores (AFS) of all methods across eight datasets. Higher values indicate better functional properties in the dataset. **B****old results** indicate the best value among all methods assessed using the same oracles.

| Methods | avGFP | AAV | TEM | E4B | AMIE | LGK | Pab1 | UBE2I | Average |
| --- | --- | --- | --- | --- | --- | --- | --- | --- | --- |
| AdaLead | 2.94 ± 0.63 | -1.88 ± 0.33 | 0.23 ± 0.19 | -0.47 ± 0.09 | -2.11 ± 0.26 | -0.07 ± 0.05 | 0.33 ± 0.30 | 2.56 ± 0.18 | 0.19 |
| DyNA PPO | 2.42 ± 0.09 | -4.31 ± 0.11 | 0.21 ± 0.03 | -1.06 ± 0.04 | -40.91 ± 0.83 | -2.71 ± 1.27 | 0.12 ± 0.00 | 1.85 ± 0.03 | -5.55 |
| CMA-ES | 2.61 ± 0.33 | -5.95 ± 0.14 | 0.19 ± 0.03 | -1.33 ± 0.07 | -38.68 ± 1.21 | -9.41 ± 0.69 | 0.11 ± 0.01 | 1.93 ± 0.06 | -6.32 |
| DbAS | 2.54 ± 0.04 | -4.70 ± 0.08 | 0.13 ± 0.01 | -1.10 ± 0.02 | -41.52 ± 0.33 | -5.08 ± 0.52 | 0.12 ± 0.00 | 1.90 ± 0.02 | -5.97 |
| CbAS | 2.54 ± 0.07 | -4.65 ± 0.07 | 0.13 ± 0.01 | -1.11 ± 0.02 | -41.24 ± 0.38 | -5.72 ± 0.21 | 0.12 ± 0.00 | 1.92 ± 0.03 | -6.00 |
| COMs | 1.75 ± 0.04 | -6.07 ± 0.03 | 0.01 ± 0.01 | -1.38 ± 0.02 | -35.92 ± 0.43 | -12.55 ± 0.38 | 0.09 ± 0.00 | 1.35 ± 0.03 | -6.59 |
| PEX | 3.14 ± 0.14 | -1.10 ± 0.45 | 0.13 ± 0.02 | 1.80 ± 1.38 | -0.91 ± 0.11 | -0.19 ± 0.05 | 0.79 ± 0.14 | 1.53 ± 0.19 | 0.65 |
| GFN-AL | 3.83 ± 0.30 | -4.57 ± 0.21 | 0.30 ± 0.02 | -1.59 ± 0.11 | -37.36 ± 1.49 | -6.57 ± 0.34 | 1.17 ± 0.05 | 3.49 ± 0.11 | -3.52 |
| GGs | 3.53 ± 0.03 | -3.09 ± 0.19 | 0.28 ± 0.05 | -1.52 ± 0.07 | -8.32 ± 0.49 | -3.12 ± 0.14 | 0.01 ± 0.00 | 1.74 ± 0.22 | -1.31 |
| MLDE | **9.33 ± 0.40** | **1.99 ± 0.02** | **2.04 ± 0.06** | **10.55 ± 0.13** | **0.03 ± 0.00** | **0.04 ± 0.01** | **1.27 ± 0.06** | **4.01 ± 0.12** | **3.67** |

#### 3) Mutations to best design

Table VI presents the number of mutations required to transform the wild-type sequence into the best-designed sequence within our final proposed population, along with the corresponding percentages relative to the actual length of the protein. Overall, our method modifies the wild-type sequences within the range of 17.9% to 45.4%, with a predominant occurrence around 31% for the majority of tasks. These figures align with natural protein diversity, as proteins within the same family exhibit variability. For instance, [82] demonstrated the existence of multiple GFP proteins in nature, with cgreGFP exhibiting the highest fluorescence despite only sharing 41% sequence similarity with the avGFP utilized in our study.

**Table VI.** Edit distance between the wild-type sequence and the best-designed sequence.

| Task | dist(WT) | Task | dist(WT) |
| --- | --- | --- | --- |
| avGFP | 107.60 ± 2.94 (45.4%) | AMIE | 124.40 ± 9.18 (36.5%) |
| AAV | 5.00 ± 0.00 (17.9%) | LGK | 149.40 ± 6.15 (34.0%) |
| TEM | 70.80 ± 2.64 (24.8%) | Pab1 | 26.00 ± 1.27 (34.7%) |
| E4B | 31.40 ± 2.42 (30.8%) | UBE2I | 42.00 ± 4.69 (26.4%) |

### F. Analysis

#### 1) Effect of masking strategy

We conduct an ablation study to demonstrate how each masking strategy contributes to the performance. In particular, we compare MLDE with its variants, including (1) without random masking (w/o RM): we do not use random masking for MLDE and (2) without importance masking (w/o IM): we only rely on random masking to select positions to mutate. As shown in the lower section of Table II, removing either component leads to a decrease in algorithm performance. This observation highlights the effectiveness of employing both strategies, affirming the efficacy of our proposed masking approach.

#### 2) Effect of beam size

The impact of varying beam sizes is depicted in Fig. 3. The graphic illustrates that increasing the beam size leads to accelerated convergence of the pipeline and results in sequences with higher fitness values throughout each iteration. This further underscores the effectiveness of incorporating the beam search technique.

**Fig. 3.**
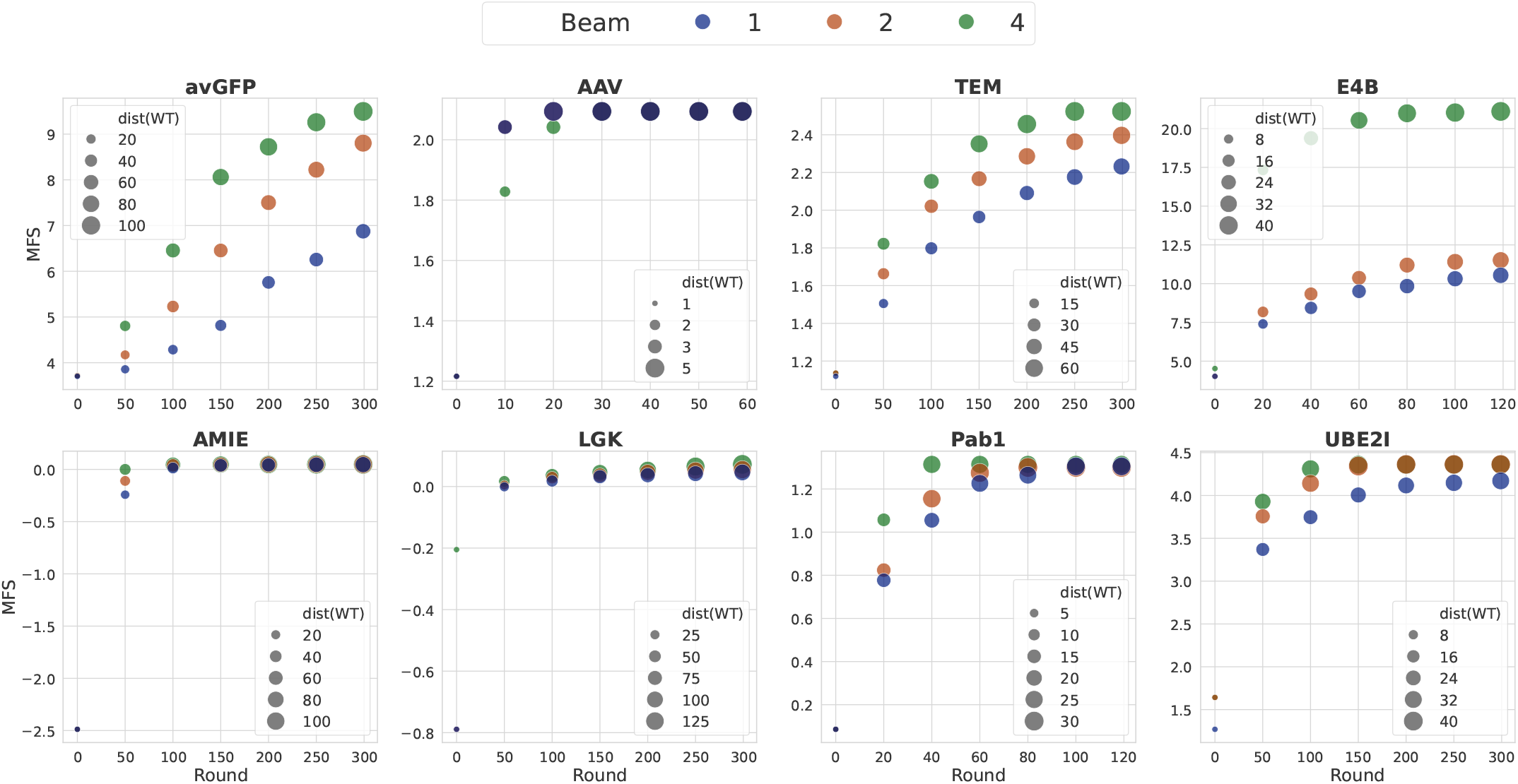
Effects of different beam sizes (k-mer size is fixed to 1) on eight tasks. The x-axis is the number of rounds, and the y-axis is the fitness score. Size of dots represents the edit distance from those particular sequences to wild-type one.

#### 3) Effect of k-mer size

The efficacy of selecting the k-mer size for sequence masking is illustrated in Fig. 4. It is evident that the influence of the k-mer size varies across datasets. For instance, while the distinction among settings is evident in the AAV dataset, with a k-mer size of 1 being the optimal setting, no significant difference is observed when we examine LGK and AMIE datasets. Therefore, it is apparent that the impact of the k-mer size fluctuates and requires careful adjustment in order to achieve optimal outcomes.

**Fig. 4.**
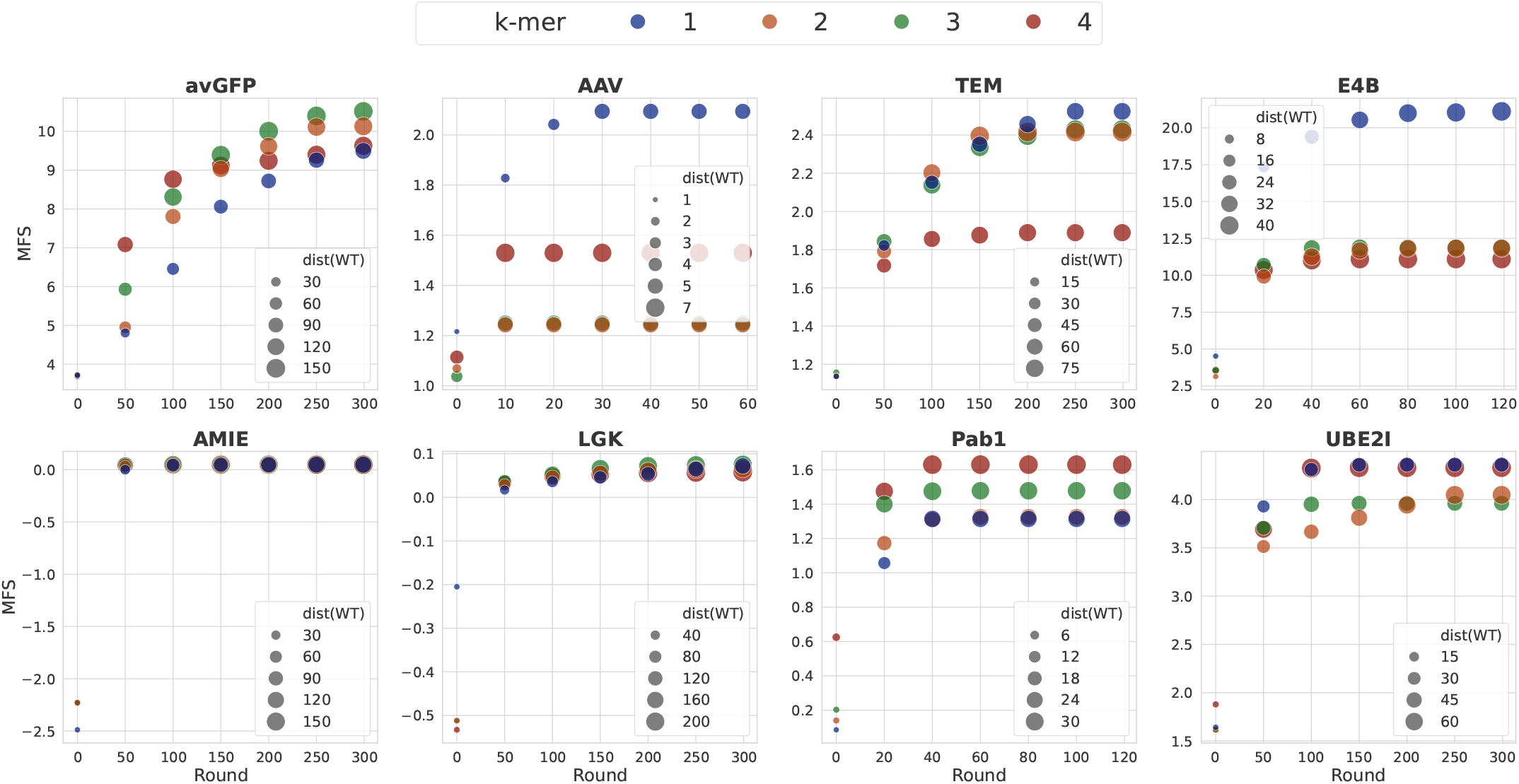
Effects of different k-mer sizes to mask (beam size is fixed to 4) on eight tasks. The x-axis is the number of rounds, and the y-axis is the fitness score. Size of dots represents the edit distance from those particular sequences to wild-type one.

#### 4) Validation of generated sequences

To comprehensively evaluate the generated samples, we assess the folding ability of proteins by using ESMFold [55]. There are three main reasons for us to decide utilizing ESMFold: (1) It generates **state-of-the-art three-dimensional structure predictions** directly from the primary protein sequence. (2) It operates without reliance on multiple-sequence alignment (MSA), a crucial but computational-burden component in the architecture of many established folding models [83], [84], resulting in a 60x speedup for short sequences and a **10x speedup on average** compared to these models. (3) It predicts structures with confidence and does not depend on any templates, **mitigating the potential for producing an “unfolded” structure** for a sequence highly similar to a template. Figure 5 illustrates the tertiary structure of TEM’s best variant, aligned with its wild-type structure. The root mean square deviation (RMSD) between the two structures is 0.324 *Å*. Additionally, the figure demonstrates the high confidence of our predicted structure, validating our model’s capability to design a real TEM-1 *β*-Lactamase protein. Structures of other proteins are presented in the Supplementary Material. We also explore the application of ESM-1v [85] in the zero-shot validation of generated sequences, which is introduced in the Supplementary Material.

**Fig. 5.**
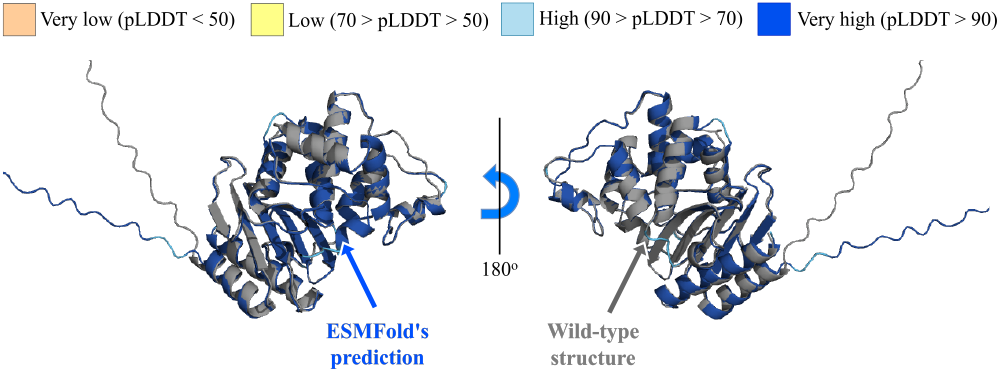
3D visualization of best-designed variant of TEM-1 *β*-Lactamase protein (colored), aligned with its wild-type structure (gray). The color of predicted structure represents ESMFold’s local prediction confidence (pLDDT) per amino acid location.

Our implementation is publicly available at https://github.com/HySonLab/Directed_Evolution.

## VI. Concluding Remarks

This paper introduces a Machine Learning-Driven Evolution (MLDE) framework. Incorporating two distinct masking strategies, namely random masking and importance masking, our pipeline utilizes pretrained models with minimal training, resulting in protein sequences exhibiting higher fitness. Experimental results across six protein sequence design tasks demonstrate that our method surpasses several robust baselines across nearly all metrics. In the future work, we aim to apply our Directed Evolution framework to the task of ligandbinding protein redesign, antibody design, etc. We hope our framework will enable scientists and researchers to efficiently optimize proteins with certain given properties.

### Limitations

Despite the positive outcomes of our approach, the protein sequences we created have not been verified through *in-vitro* experimentation, which introduces a level of uncertainty associated with the oracle model. The results presented in our research are obtained by averaging data from five separate runs with different seeds, which can accommodate some variance on these metrics. Additionally, all fitness data utilized in our study are derived from wetlab experiments, ensuring that the fitness values within the training datasets are realistic. Although there is currently no perfect metric available, we believe that new methods should be encouraged in the field of computational biology.

