## Appendix for "Protein Design by Directed Evolution Guided by Large Language Models"

### I. Data Descriptions

*1) Green Fluorescent Proteins (**avGFP**):* Derived from Aequorea victoria, Green Fluorescent proteins (GFPs) are capable of manifesting vivid green fluorescence upon exposure to light within the blue to ultraviolet spectrum. These proteins are commonly employed as biosensors for detecting gene expressions and protein locations. Our objective is to design sequences with enhanced capabilities as gene delivery vectors, quantified by AAV viabilities. The magnitude of the search space encompasses $20^{28}$ possibilities.

*2) Adeno-associated Viruses (**AAV**):* The engineer of a 28-amino acid segment (position 561–588) within the VP1 protein, situated in the capsid of the Adeno-associated virus, has garnered significant interest in the realm of machine learning-guided design. We aim to design more capable sequences as gene delivery vectors measured by AAV viabilities. The size of the search space is $20^{28}$.

*3) Aliphatic Amide Hydrolase (**AMIE**):* The enzyme encoded by amiE, known as Amidase, holds industrial relevance and is d erived from Pseudomonas aeruginosa. Our objective is to optimize amidase sequences that result in enhanced enzyme activities, defining a search space comprising $20^{341}$ sequences.

*4) Ubiquitination Factor Ube4b (**E4B**):* The ubiquitination factor Ube4b plays a pivotal role in cellular waste degradation through interactions with ubiquitin and other proteins. Our emphasis is on the enhancement of E4B enzyme activity within the specified landscape, which encompasses a search space of $20^{102}$.

*5) Levoglucosan Kinase (**LGK**):* Levoglucosan kinase catalyzes the conversion of Levoglucosan (LG) to the glycolytic intermediate glucose-6-phosphate through an ATP-dependent reaction. The target is to optimize LGK protein sequences with improved enzyme activity. The size of search space is $20^{439}$.

*6) Poly(A)-binding Protein (**Pab1**):* Pab1 utilizes the RNA recognition motif (RRM) to bind to multiple adenosine monophosphates (poly-A). The aim is to design sequences with higher binding fitness to multiple adenosine monophosphates. The search space size is $20^{75}$ on a segment of the wild-type sequence.

*7) TEM-1 $\beta$-Lactamase (**TEM**):* The investigation of the TEM-1 $\beta$-Lactamase protein's resistance to penicillin antibiotics in E. coli is a subject of extensive scrutiny, aiming to comprehend mutational impacts and the associated fitness landscape. The optimization objective involves suggesting sequences with elevated thermodynamic stability compared to the wild-type TEM-1 within a search space of magnitude $20^{286}$.

*8) SUMO E2 conjugase (**UBE2I**):* The utilization of variants for the functional mapping of human genomes holds considerable significance in both scientific research and clinical treatment. The search space encompasses a size of $20^{159}$ within a segment of the wild-type sequence.

### II. Evaluation Metrics

We provide mathematically definitions of five metrics: maximum fitness score (MFS), average fitness score (AFS), diversity, novelty, and dist(WT). Let $x^{wt}$ be the wild-type sequence, $P = \{p_1, p_2, \ldots, p_N\}$ be the population generated by MLDE, and $\mathcal{O}(\cdot)$ be the oracle predicting the fitness score of a sequence, we define:

- **MFS** = $\max(\{\mathcal{O}(p_i)\}_{i=1}^{N})$,
- **AFS** = $\dfrac{\sum_{i=1}^{N} \mathcal{O}(p_i)}{N}$
- **Diversity** = $\dfrac{\sum_{i=1}^{N} \sum_{j=1, j\neq i}^{N} d(p_i, p_j)}{N(N-1)}$,
- **Novelty** = $\dfrac{\sum_{i=1}^{N} \min_{s_j \in \mathcal{D}} d(p_i, s_j)}{N}$,
- **dist(WT)** = $d(\arg\max_{p_i \in P} \mathcal{O}(p_i), x^{wt})$

where $d(\cdot, \cdot)$ is the Levenshtein distance (i.e., edit distance), and $\mathcal{D}$ is the initial dataset (i.e., training dataset).

### III. Oracles

We establish the optimization oracle by leveraging features generated by the pre-trained 33-layer ESM-2 [1] with a dimension of 1280. Subsequently, we train an Attention1D module to predict fitness values based on these representations. As for the evaluation oracle, which acts as a "ground-truth" evaluator, we employ the trained oracle provided by [2]. This

Corresponding author: Truong Son Hy.
Truong Son Hy is with University of Alabama at Birmingham, United States
Both authors are with FPT Software AI Center, Hanoi, Vietnam

TABLE I: Masked marginal probability of generated sequences on eight datasets.

| Methods | avGFP | AAV | TEM | E4B | AMIE | LGK | Pab1 | UBE2I | Pearson $r$ |
|---|---|---|---|---|---|---|---|---|---|
| PEX | -4460.32 | -493.39 | -5161.91 | -1860.53 | -6141.57 | -7854.06 | -1357.85 | -2843.76 | 0.61 |
| GFN-AL | **14.82** | **0.05** | -20.97 | **0.82** | **10.49** | **12.83** | **-0.31** | -10.11 | -0.35 |
| MLDE | -2.46 | 0.01 | **-0.02** | 0.02 | -6.08 | -0.05 | -0.63 | **2.21** | 0.19 |
| Total | | | | | | | | | -0.05 |

TABLE II: Metrics across eight benchmarks when utilizing ESM-1v as the oracle.

| Metric | avGFP | AAV | TEM | E4B | AMIE | LGK | Pab1 | UBE2I |
|---|---|---|---|---|---|---|---|---|
| Fitness$^{(\mathrm{WT})}$ | **3.68** | **-2.73** | **1.08** | **0.77** | **-2.79** | **-1.26** | 0.01 | 0.77 |
| MFS | 3.50 | -4.80 | 0.01 | -1.06 | -11.99 | -3.68 | **0.02** | **2.00** |
| $\mathcal{S}_m$ | 81.23 | 9.14 | 72.98 | 17.73 | 68.56 | 88.77 | 14.72 | 53.82 |

evaluation oracle combines the pre-trained ESM-1b [3] with an Attention1D module with a dimension of 512, and it is used to assess all methods, including both baselines and our proposed approach.

### IV. Baselines' Hyperparameters Selection

This section outlines how we choose the hyperparameters of the baselines. Among the nine baselines used for comparison, we employ the released code and default settings for PEX, GFN-AL, and GGS, as these baselines have been developed, benchmarked, and hyperparameters tuned on protein datasets. For AdaLead, the primary tunable hyperparameters are the recombination rate $r$, mutation rate $\mu$, and the threshold $k$. While the algorithm has been demonstrated to be fairly robust in performance for $k, r < 0.5$ [4], we found that $\mu$ is task-specific with an optimal range of $[1, 3]$. For DyNA PPO, the primary tunable hyperparameter is the score threshold $\tau$, which was found to be optimal at $\tau = 0.5$ in the original paper [5]. For the remaining baselines, we follow the hyperparameter selection workflow proposed by [6] to determine appropriate hyperparameters for each method.

### V. Validating the Generated Sequences Using ESMFold

Continuing the analysis from the main text, this section demonstrates that our method can generate high-fitness and foldable protein sequences. We begin by utilizing ESMFold [1] to fold wild-type sequences, focusing only on datasets with outputs displaying high confidence. This approach ensures that we test reliable structures for mutants by verifying the reliability of their corresponding wild-type structures. Figure 1a, 1b, 1c, and 1d illustrate the predicted 3D structures of best-designed sequences of E4B, AMIE, Pab1, and UBE2I, respectively. From these figures, it is evident that there exists a significant degree of overlap between our designed sequences and the wild-type proteins. These empirical findings presented in our study suggest that our method can produce genuine proteins. While further investigation is required to determine if the suggested sequences can effectively expedite wet-lab studies, these *in-silico* results can be seen as encouraging indicators for the field of protein design, affirming the effectiveness of our method.

### VI. Bonus 1: On the Optimization Performance with Zero-Shot Prediction of Mutational Effects

Despite the effectiveness of current approach, our method, as well as many other deep learning-based approaches, still require a small amount of training data for the specific type of protein being optimized. This data is typically obtained through deep mutational scanning, which is costly and time-intensive. ESM-1v [7] offers a potential solution to this challenge by serving as a protein language model capable of performing *zero-shot* prediction of mutational effects across various proteins with diverse functions. This capability enables the estimation of the log-likelihood of generated samples relative to the wild-type sequences without the need for additional training or collecting experimental assay data. Specifically, let $x^{mt}$ and $x^{wt}$ denote the mutant and wild-type sequences, respectively. We denote $x_{-M}$ as the sequence $x$ with a mask introduced at a set of positions $M$. The score for a mutation is computed by considering its probability relative to the wild-type amino acids, known as the masked marginal probability:

$$\mathcal{S}_m = \sum_{i \in M} \log p(x_i = x_i^{mt} | x_{-M}) - \log p(x_i = x_i^{wt} | x_{-M}) \tag{1}$$

While it cannot directly predict the fitness value of sequences, this model can validate whether the generated samples are plausible in terms of structure and function, serving as an indirect measure of fitness. We hypothesize that this model can replace the Attention1D module, the only part requiring additional training, to optimize sequences. To test this hypothesis, we conduct two additional experiments: (1) We employ ESM-1v to validate the best-designed sequences produced by our current method and most recent baselines. (2) We utilize ESM-1v as the optimization oracle to design new proteins and validate the generated samples using the evaluation oracle described in Section III.

Table I outlines the masked marginal probability (Equation (1)) of generated sequences across eight datasets and their correlation with the reported fitness values. It is evident that there is no clear relationship between these metrics, as the Pearson correlation drastically varies among methods. When combining all of these values, the correlation approaches zero, indicating a lack of relationship between these two metrics.

TABLE III: Maximum fitness score (MFS) of proposed method with different pLMs. Higher values indicate better functional properties in the dataset. **Bold** results indicate the best value, and underlined results indicate the second-best value.

| Input type | pLM | avGFP | AAV | TEM | E4B | AMIE | LGK | Pab1 | UBE2I |
|---|---|---|---|---|---|---|---|---|---|
| Residue only | ESM-2 | 9.50 ± 0.37 | 2.09 ± 0.00 | 2.54 ± 0.06 | **11.56 ± 0.14** | **0.05 ± 0.00** | **0.07 ± 0.01** | 1.32 ± 0.06 | **4.41 ± 0.12** |
| | SaProt w/ seqOnly | **9.71 ± 0.79** | 2.13 ± 0.03 | 2.60 ± 0.08 | 10.24 ± 0.42 | **0.05 ± 0.00** | 0.06 ± 0.00 | 1.51 ± 0.02 | 3.96 ± 0.02 |
| Structure-aware | SaProt | 6.74 ± 0.64 | **4.07 ± 0.22** | **2.82 ± 0.10** | 10.42 ± 0.31 | **0.05 ± 0.00** | 0.04 ± 0.00 | **1.60 ± 0.12** | 3.94 ± 0.05 |

In the second experiment, Table II shows the fitness values achieved when using ESM-1v as an optimization oracle. It can be observed that while the masked marginal scores are high, the model fails to optimize sequences, as their fitness values decrease dramatically, with the majority even falling below their wild-type fitness. These findings, once again, indicate that we cannot directly utilize ESM-1v as the oracle for optimizing protein sequences.

### VII. Bonus 2: On the Impact of Large Language Models To the Overall Performance

Advancements in deep neural network architectures tailored for sequences, along with pretraining on extensive datasets, have revolutionized automated text analysis [8], [9]. Inspired by these developments, many pre-trained Protein Language Models (pLMs) have emerged, becoming essential tools for protein-related tasks in bioinformatics. In this section, we investigate whether utilizing more advanced pLMs leads to improved optimization results. Specifically, we compare the 35M-parameter **ESM-2** with the 35M-parameter **SaProt** [10], which is the state-of-the-art model in the ProteinGym benchmark [11]. While SaProt is a structure-aware pLM, it also has a *sequence-only* version (**SaProt w/ seqOnly**), which does not incorporate structural information into the input sequences. For more detailed information, readers are encouraged to refer to the original paper, as this is outside the scope of our study. Since our benchmark datasets do not include structural information for wild-type proteins, we use AlphaFold 3 [12] to predict the tertiary structure of each protein[1], which we then use to generate structure-aware sequences. Table III presents the comparison results when these pLMs are integrated into the proposed pipeline. It is observed that SaProt, with the same number of parameters, significantly impacts overall performance, increasing the MFS in 4 out of 8 tasks. In the AAV task, it nearly doubles the performance (4.07 for SaProt compared to 2.09 for ESM-2). Even when considering second-place finishes, SaProt w/ seqOnly still outperforms ESM-2, winning in every tasks where structure-aware SaProt excels. These results empirically demonstrate that the emergence of pLMs can enhance the performance of our proposed method. This is a positive outcome, indicating that our work can benefit from advancements in the protein design field.

Our implementation is publicly available at https://github.com/HySonLab/Directed_Evolution.

[1]We do not use ESMFold since it has a known issue with GFP structure prediction.

[10] J. Su, C. Han, Y. Zhou, J. Shan, X. Zhou, and F. Yuan, "Saprot: Protein language modeling with structure-aware vocabulary," in *The Twelfth*

*International Conference on Learning Representations*, 2024. [Online]. Available: https://openreview.net/forum?id=6MRm3G4NiU

[11] P. Notin, A. Kollasch, D. Ritter, L. Van Niekerk, S. Paul, H. Spinner, N. Rollins, A. Shaw, R. Orenbuch, R. Weitzman *et al.*, "Proteingym: large-scale benchmarks for protein fitness prediction and design," *Advances in Neural Information Processing Systems*, vol. 36, 2024.

[12] J. Abramson, J. Adler, J. Dunger, R. Evans, T. Green, A. Pritzel, O. Ronneberger, L. Willmore, A. J. Ballard, J. Bambrick, S. W. Bodenstein, D. A. Evans, C.-C. Hung, M. O'Neill, D. Reiman, K. Tunyasuvunakool, Z. Wu, A. Žemgulytė, E. Arvaniti, C. Beattie, O. Bertolli, A. Bridgland, A. Cherepanov, M. Congreve, A. I. Cowen-Rivers, A. Cowie, M. Figurnov, F. B. Fuchs, H. Gladman, R. Jain, Y. A. Khan, C. M. R. Low, K. Perlin, A. Potapenko, P. Savy, S. Singh, A. Stecula, A. Thillaisundaram, C. Tong, S. Yakneen, E. D. Zhong, M. Zielinski, A. Žídek, V. Bapst, P. Kohli, M. Jaderberg, D. Hassabis, and J. M. Jumper, "Accurate structure prediction of biomolecular interactions with alphafold 3," *Nature*, May 2024. [Online]. Available: http://dx.doi.org/10.1038/s41586-024-07487-w

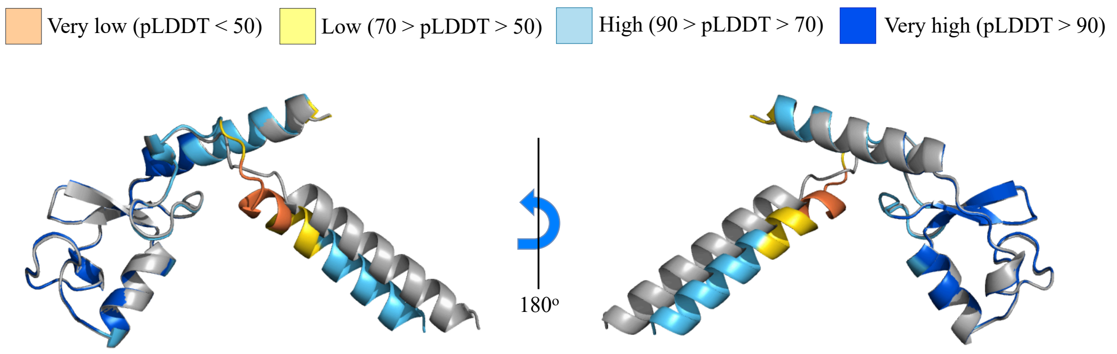


(a) E4B structures with RMSD = 0.287$\mathring{A}$

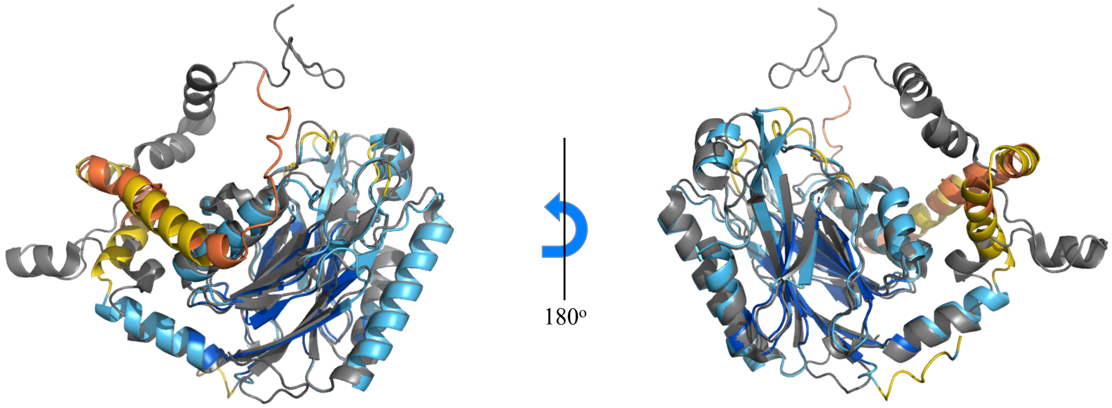


(b) AMIE structures with RMSD = 1.148$\mathring{A}$

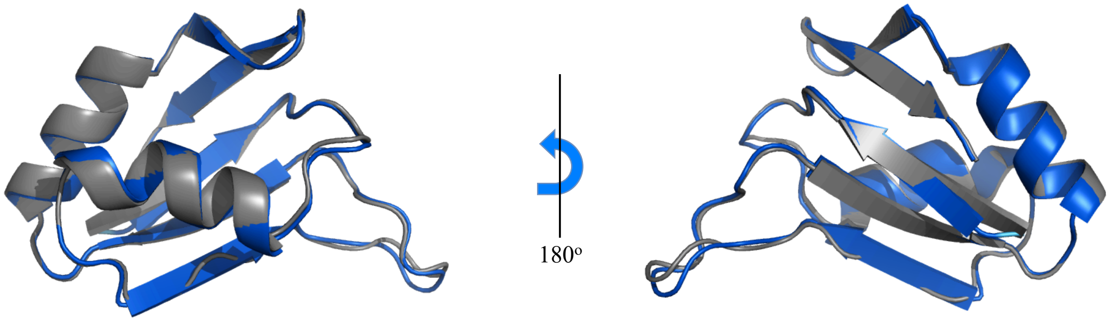


(c) Pab1 structures with RMSD = 0.38$\mathring{A}$

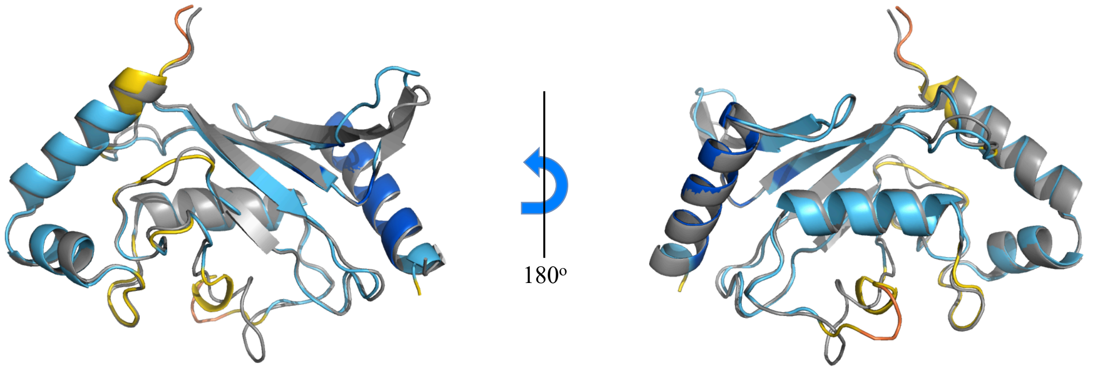


(d) UBE2I structures with RMSD = 0.528$\mathring{A}$

Fig. 1: 3D visualizations of best-designed sequences generated by our method (colored), aligned with their corresponding wild-type structure (gray). The color of predicted structure represents ESMFold's local prediction confidence (pLDDT) per amino acid location.
